# Raynals, an online tool for the analysis of dynamic light scattering

**DOI:** 10.1101/2023.04.25.538274

**Authors:** Osvaldo Burastero, George Draper-Barr, Bertrand Raynal, Maelenn Chevreuil, Patrick England, Maria M. Garcia-Alai

## Abstract

Dynamic light scattering (DLS) is routinely employed to assess the homogeneity and size distribution profile of samples containing microscopic particles in suspension or solubilised polymers. In this work, we introduce Raynals, an user-friendly software for the analysis of single-angle DLS data that uses the Tikhonov-Phillips regularisation. Performance is evaluated on simulated and experimental data, generated by different DLS instruments, for several proteins and gold nanoparticles. DLS data can be easily misinterpreted and the simulation tools available in Raynals allow understanding of the limitations of the measurement and its resolution. It has been designed as a tool to address quality control of biological samples, during sample preparation and optimisation, and it helps in the detection of aggregates showing the influence of large particles. Last, Raynals provides flexibility in the way the data is presented, allows exporting publication-quality figures, it is free for academic use, and can be accessed online on the eSPC data analysis platform at spc.embl-hamburg.de.

## 1. Introduction

The polarisability of light by biological samples is not homogeneous due to macromolecules in suspension undergoing Brownian motion and therefore an incident light beam scatters in all directions and the scattering intensity fluctuates over time. From the total intensity of the scattered light, valuable information about the molecular weight, radius of gyration, internal spatial arrangement of scattering centres, and virial coefficients can be obtained (Schmitz & Phillies, 1991). Moreover, the temporal behaviour of the scattered light allows the estimation of an apparent translational diffusion coefficient D_app_, and its related hydrodynamic radius (R_h_) (Schmitz & Phillies, 1991).

Dynamic light scattering (DLS) experiments consist of shooting a polarised monochromatic laser into a sample at short time intervals. The diffracted light undergoes constructive or destructive interference by the surrounding particles generating an intensity fluctuation that correlates to the time scale movements of the particles. This dynamic information of the particles scattering is collected after passing through a second polarizer. The second-order autocorrelation function is then constructed from the acquired intensity trace to determine D_app_. Samples that can be fitted using a single exponential decay are considered to be monodisperse. In addition, polydisperse systems would require a sum of exponential decays. D_app_ can be estimated at single or multiple angles, with angular dependence of the signal revealing scattering particle shape.

DLS is a fast and low-consumption method routinely used to assess the homogeneity and aggregation state of protein samples (Raynal *et al*., 2014; de Marco *et al*., 2021; Stetefeld *et al*., 2016). It can also be used for more advanced applications: such as the calculation of critical micelle concentrations (CMC) of detergents (Sutherland *et al*., 2009), optimisation of solutions for protein-detergent complexes (Meyer *et al*., 2015), and to study protein crystallisation (Saridakis *et al*., 2002; Dierks *et al*., 2008; Meyer *et al*., 2012; Oberthuer *et al*., 2012; Schubert *et al*., 2017). A limitation of this technique is that the scattering signal is extremely sensitive to the particle ratios, so large aggregates overshadow the signal from smaller particles, even if the latter population is a greater proportion of the total particle number in solution. In addition, the correlogram deconvolution is an ill-posed problem implying that is not possible to retrieve the original intensity-weighted particle size distribution. For single-angle measurements, the rule of thumb is that only species that differ by a factor of two/three in their R_h_ can be totally distinguished, and technical instrumentation limitations could expand this factor to a range of five. Finally, the transformation of the intensity distribution to mass (or volume) distribution is subject to additional assumptions such as assuming that particles are perfect hard spheres with constant density.

Results from DLS experiments are typically analysed using the commercial software provided by the instrument vendor. Occasionally, advanced users decide to fit their data with desktop programs like CONTIN or SEDFIT (Provencher, 1982; Brown *et al*., 2007). Here, we introduce Raynals, an online tool designed for the interpretation of DLS data tailored for biological samples. Raynals is our newest addition to the eSPC (spc.embl-hamburg.de), an online data analysis platform that contains, so far, modules for evaluating experimental data from differential scanning fluorimetry, mass photometry, and microscale thermophoresis (Burastero *et al*., 2021; Niebling *et al*., 2021, 2022).

## 2. Results and Discussion

### 2.1. Raynals workflow

The Raynals tool usage workflow can be divided into four steps (Figure 1). To begin with, the user loads the normalised second-order autocorrelation curves and the associated experimental information, such as the angle of detection and laser wavelength. Raw curves can be filtered by removing those with a lower intercept or a “bumpy” baseline to exclude samples with aggregates and/or buffers. Sample preparation by centrifugation and filtration is a critical step to remove dust particles and aggregates from the solution that would introduce artefacts to the measurements.

**Figure 1.**
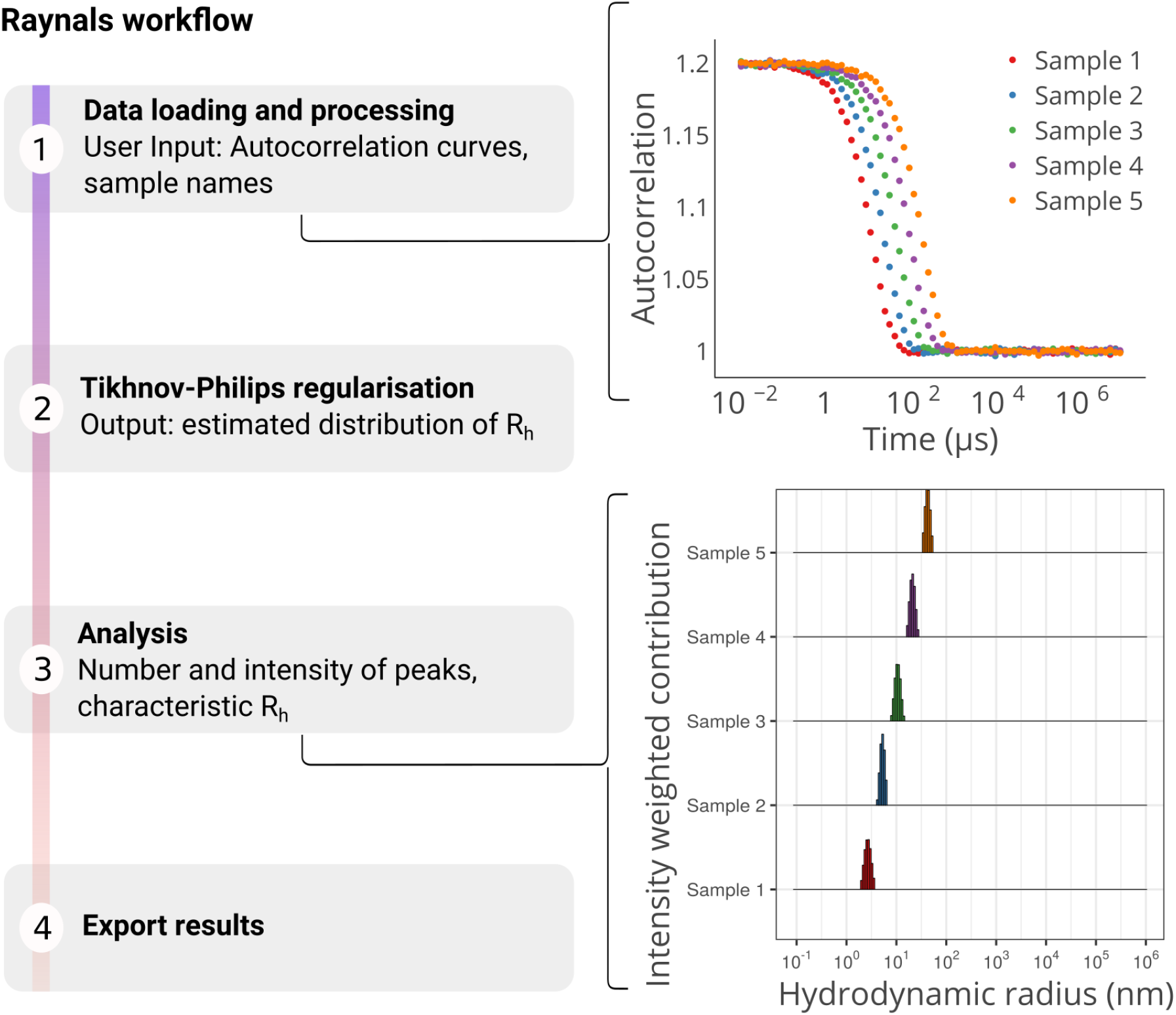
The Raynals pipeline has four steps. Firstly, the second-order autocorrelation data is loaded and filtered based on initial value and baseline quality. Input parameters include detection angle, laser wavelength, temperature, refractive index, and viscosity. Secondly, a regularisation approach is used to fit the first-order autocorrelation data, assuming a smooth non-parametric distribution of decay rates (or hydrodynamic radii). Thirdly, a threshold based on residuals can be used to remove poorly fitted curves and the estimated distributions are displayed. The user must select regions of interest to extract information about the peaks (e.g., contribution to the total intensity). Finally, the user can export the R_h_ distribution and the associated second-order autocorrelation curves.

DLS data is fitted using models based on the Siegert relationship, which connects the normalised second-order g^(2)^(*τ*) and first-order g^(1)^(*τ*) autocorrelation functions. *g*^(1)^(*τ*) can be represented by an intensity-weighted integral over a distribution of decay rates G(┌) (Xu, 2006):

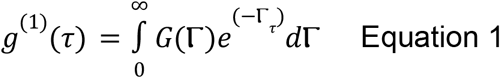

where G(┌) is normalised as follows

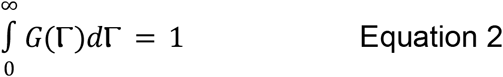

Finding the distribution G(┌) from noisy data is challenging, and there are several available methods that can be used (Schmitz & Phillies, 1991). One common approach is to use the cumulants method proposed by Koppel to estimate the mean and variance of the distribution (Koppel, 1972). These values are used to calculate the polydispersity index (PdI) and %polydispersity (%PdI).

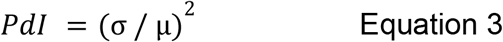

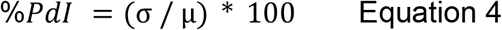

In this method, μ and σ are respectively the estimated sample mean and standard deviation. The formula for the %PdI is exactly the same as for the percentage of coefficient of variation (%CV). It has been stated that values of PdI below 0.05 (%PdI < 22) correspond to monodisperse colloidal particles; and values close to the unity (or above) indicate polydisperse samples. Despite being recommended by the International Organization for Standardization (ISO 13321 and ISO 22412), the cumulants method is highly sensitive to small amounts of aggregates and may yield misleading results in non-monodisperse samples (Mailer *et al*., 2015).

Alternative methods to determine G(┌) include adjusting a discrete number of exponentials with different decay rates that follow or not a parametric distribution. In Raynals, we implemented the fitting of *g*^(1)^(*τ*) through the Tikhonov-Phillips regularised inversion, which is commonly used to solve ill-posed inverse problems. (Phillips, 1962; Provencher, 1982; Brown *et al*., 2007). This method requires the selection of a regularisation matrix (L) and a regularisation parameter (α), and then finding the vector of relative contributions (x), such that

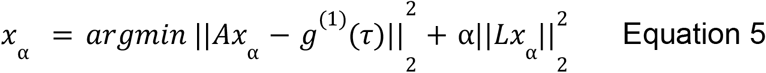

where A is the kernel matrix with values *a*_*i,j*_,

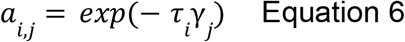

where *i* and *j* iterate over the lag time and decay rate vectors, respectively. In Raynals, the decay rate space is discretized in such a way that the hydrodynamic radius points are evenly spaced in a log scale, and L is the second-order derivative matrix that constrains how close the value of each decay rate is from its neighbours.

Thirdly, the distribution of decay rates is transformed into a distribution of diffusion coefficients and subsequently into a distribution of R_h_ (Stokes-Einstein relation, Equation 12). Then, the fitted curves can be filtered based on the residuals. To report the R_h_ (or diffusion coefficients), we apply a peak searching algorithm within user-selected intervals (e.g.1 to 100 nm). In the following sections, we assess the performance of the developed software, highlighting capabilities and limitations after analysing experimental and simulated data.

### 2.2. Addressing Raynals performance with simulated DLS data

DLS data has been simulated based on the following protocol (Figure 2). To start with, we created samples with a number-weighted gaussian distribution of R_h_. Then, using the Mie Theory, we calculated the scattered light intensity of the particles, assuming they were perfect spheres. Finally, we discretized the R_h_ space, computed the relative contributions to the total intensity of each interval, and obtained the autocorrelation curves. It’s important to note that both the intensity and number distribution are just different physical representations of the same reality. All data was generated using the ‘Simulation’ panel from Raynals. Explanatory files containing the used parameters to reproduce the simulations are provided as Supplementary Material.

**Figure 2.**
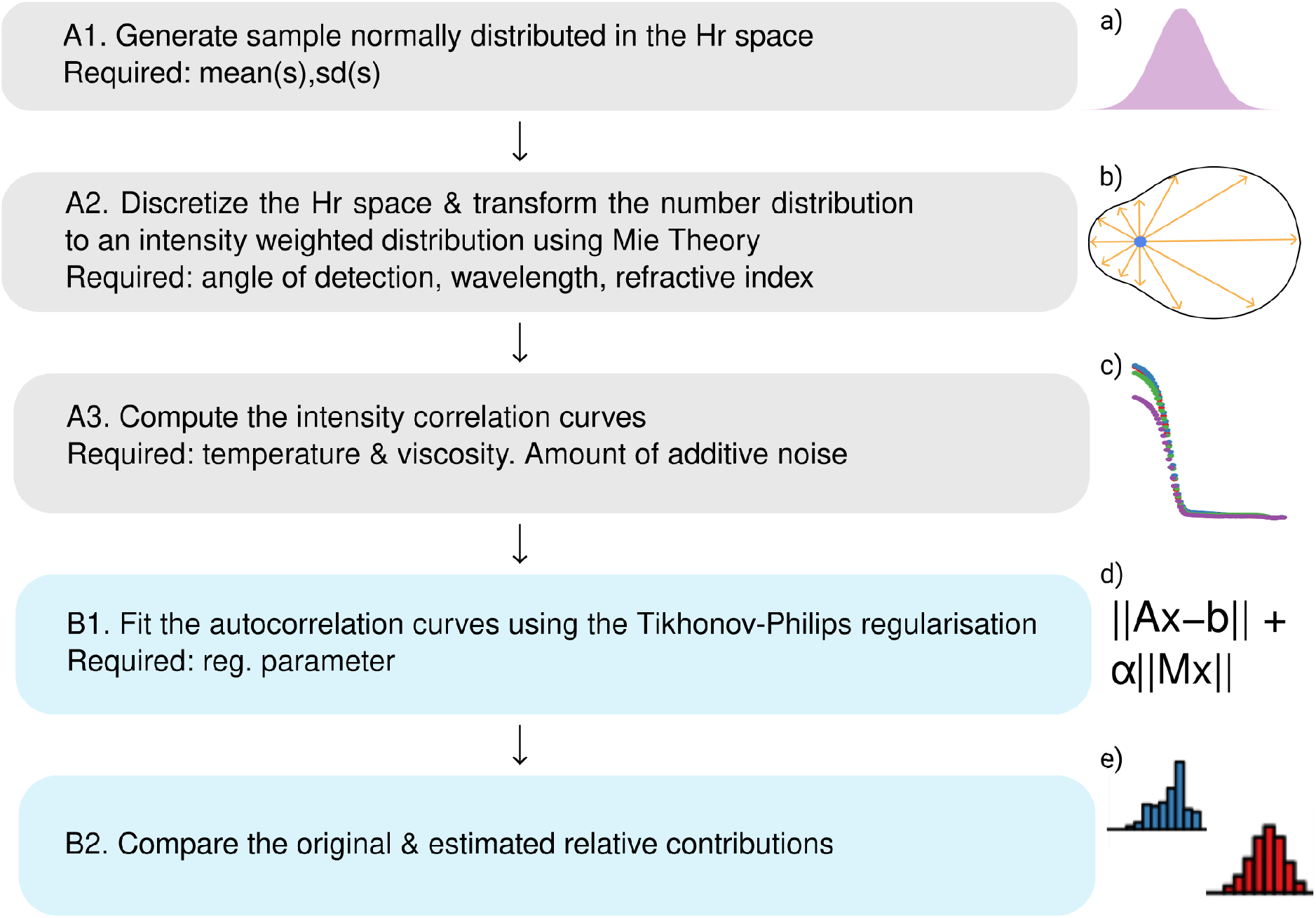
Workflow of the simulations done to evaluate the capacity of the Tikhonov-Philips regularisation to return the original hydrodynamic radius distribution. Gray and blue boxes represent respectively the data generating and data fitting steps. Figures from the right from top to bottom: a) normal distribution, b) light scattered by a particle at different angles according to the Mie Theory, c) simulated autocorrelation curves, d) equation of the regularisation approach required to solve a nonlinear inverse problem, and e) histograms of the fitted intensity distributions.

#### Case 1. One population, low %CV

To test the simplest case, we have simulated autocorrelation curves of a monodisperse sample (defined by a %CV = 10 %). It has been reported that using the weighted harmonic mean (WHM) for estimating particle size would be a better alternative than the weighted mean (WM) because it matches the average size from the cumulants analysis for low PdI samples (Farkas & Kramar, 2021). For datasets containing non-negative values, the harmonic mean is lower or equal to the geometric mean, and the geometric mean is lower or equal to the arithmetic mean (Bullen, 2003). In our analysis, we have compared the usage of WHM or the highest peak value from the distribution (mode) for obtaining R_h_. The fitted distributions are available for download in Raynals and users can choose their preferred R_h_ reporting method.

Our results show that the estimated and original R_h_ values were in excellent agreement (Figure 3) with the mean R_h_ from the number distribution remaining constant after transforming to an intensity-weighted distribution (Figure S1A). Consistently,, we observed no significant differences between using the peak maximum or the weighted harmonic mean (WHM) to estimate R_h_ (Figure S1B).

**Figure 3.**
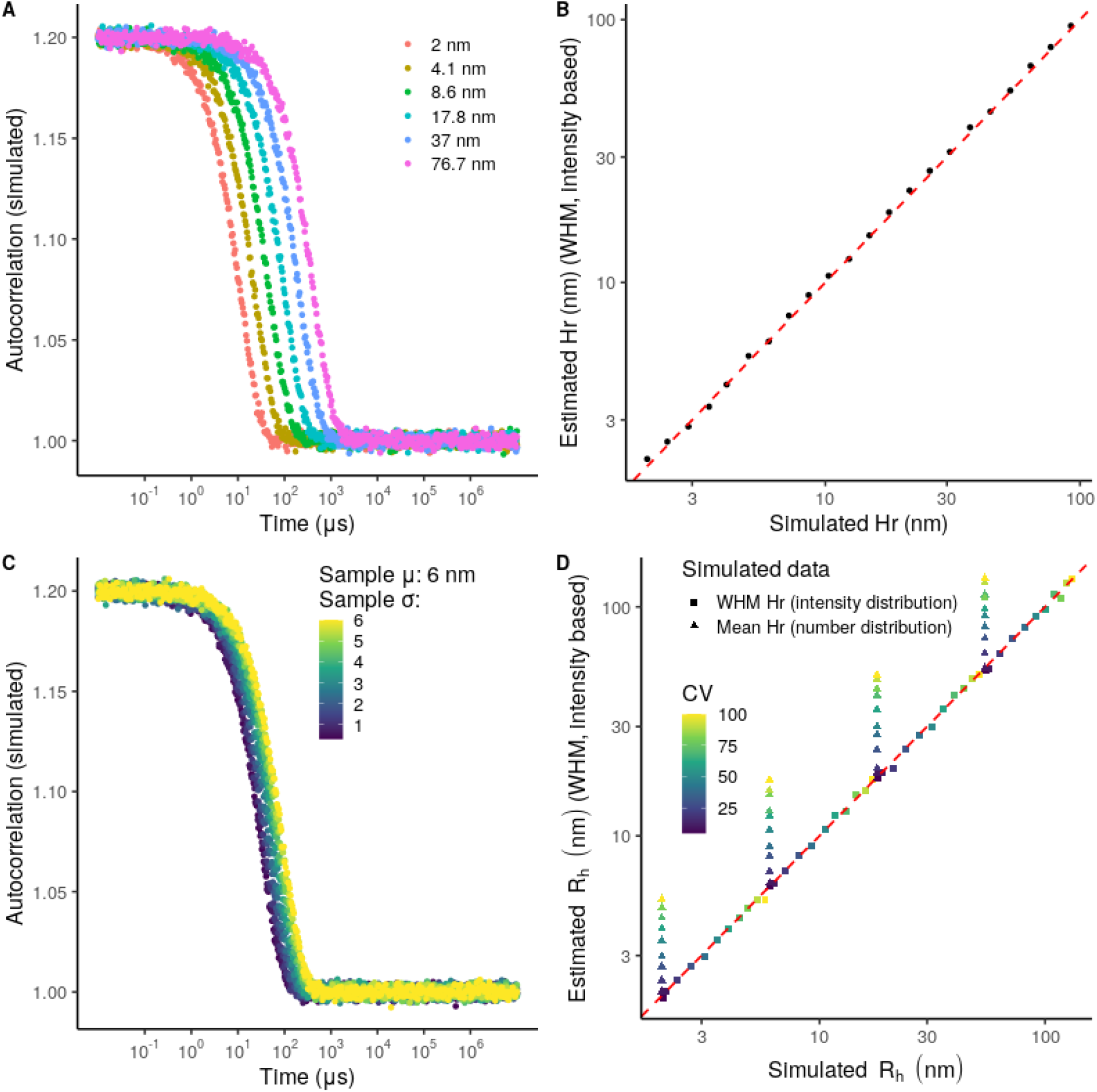
A) Simulated autocorrelation curves for samples following a number-weighted normal distribution with a %CV of 10 % (e.g., a mean of 2 nm and an sd of 0.2 nm). B) Estimated R_h_ (WHM, intensity-based) versus the mean R_h_ from the simulated number-based distributions. C) Example curves of the generated samples following a number-weighted normal distribution with a mean of 6 nm and %CVs ranging from 5 to 100 %. D) Estimated R_h_ (WHM, intensity-based) versus the WHM and mean R_h_ from the simulated intensity-based and number-based distributions, respectively. B) and D) The red line would indicate a perfect fitting. To estimate the WHM an arbitrary α of 0.01 was used.

**Figure 4.**
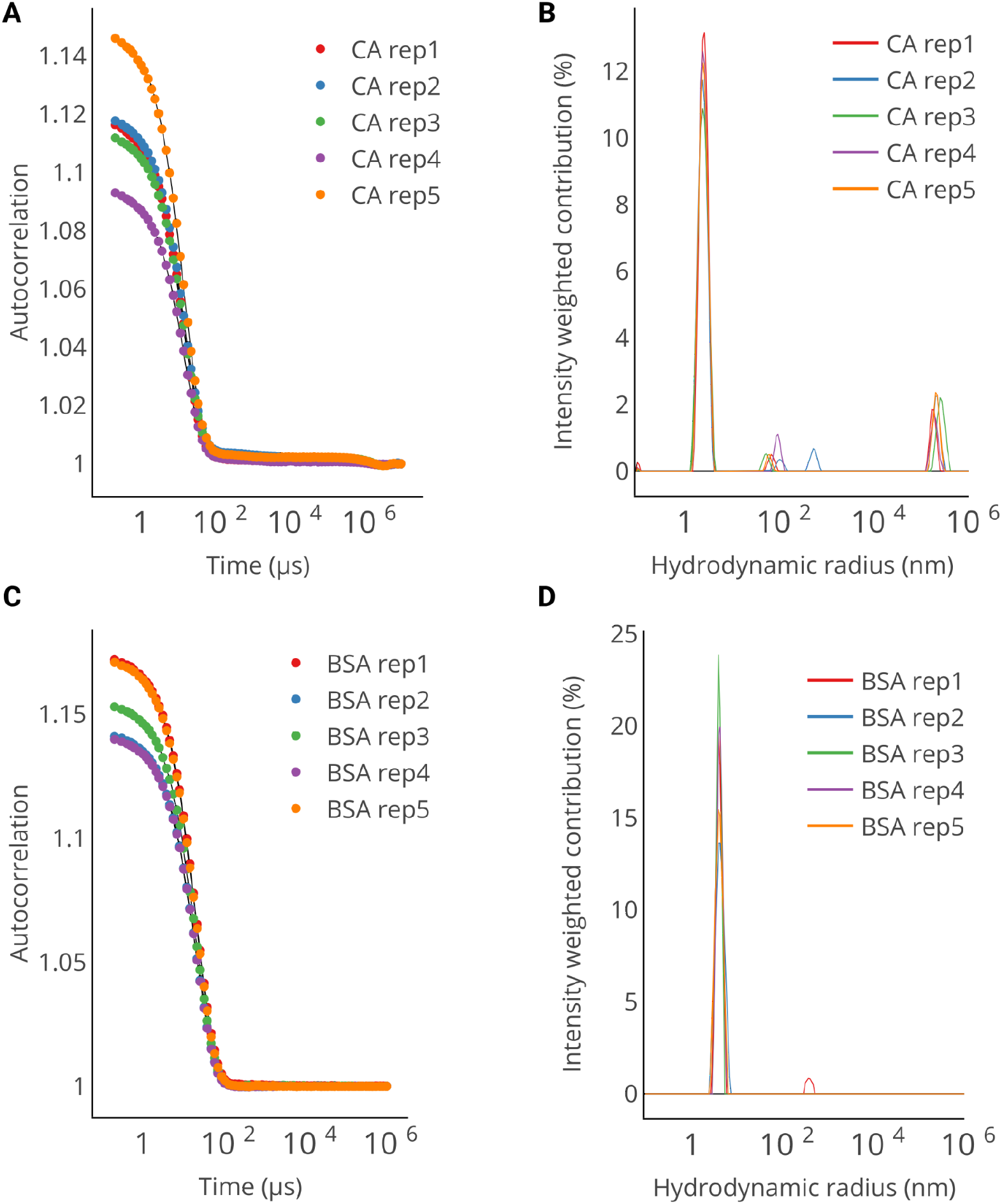
A) and C) Autocorrelation curves of carbonic anhydrase (CA) and bovine serum albumin (BSA). B) and D) Estimated relative contributions of each hydrodynamic radius for CA and BSA. The regularisation parameters were determined with the L-curve criterion.

#### Case 2. One population, increased %CV

To further evaluate the efficacy of the Tikhonov Philips regularisation method with a second-order derivative matrix as a penalisation term, we fitted 44 samples (μ = 2, 6, 18, or 54 nm) with %CVs ranging from 5 to 100 % (Figure 3C). The resulting WHM based on the fitted intensity distribution consistently correlated with the WHM derived from the simulated intensity distribution (Figure 3D, squares). However, for the samples with higher %CVs (> 25 %), the estimated WHM R_h_ did not correlate with the mean R_h_ from the simulated number distribution (Figure 3D, triangles). As expected, these simulations highlight how the algorithm can effectively retrieve the correct R_h_ from the underlying intensity distribution, but it may not be as accurate when comparing to the number distribution (Figure 3D, triangles).

It would be highly beneficial to determine whether we can recover the original particle size distribution, in addition to the “characteristic” R_h_. In this sense, the regularisation parameter α is crucial and the solution may be completely under or over-smoothed. When the amount of noise is unknown, the value of α is typically selected through an a posteriori empirical rule. To date, there is no agreed-upon criterion on how to select the mentioned rule. Some applied criteria are the L-curve (Lawson & Hanson, 1974), the U-curve (Krawczyk-Stańdo & Rudnicki, 2007), the Composite Residual and Smooth Operator (CRESO)(Cheng *et al*., 2003), the Product (Lian *et al*.), the Zero-crossing (Cheng *et al*., 2003) and the General-Cross Validation (Hansen, 1994) methods. An appropriate evaluation of the available different methods remains to be an active area of DLS research. Two questions arise naturally: 1) Can we use a fixed α for comparing the sample width? 2) Is there a way to find the optimal value of α producing an estimated distribution similar to the original one? The goal is to find a correlation between the “true standard deviation” (a proxy for polydispersity) from the generated and the estimated distributions. To address the first question, we analysed the created samples using different values of α (0.0001, 0.001, 0.01, 0.1 and 1).

Table 1 shows the correlation values between the estimated and true standard deviation values for the simulated distributions (Figure S2). These results suggest that for one population with the given level of noise, the proposed fitting method is useful for comparing standard deviations. An example of the fitted versus simulated intensity distributions is provided in Figure S3. The value of α that gives on average the closest distribution is α=1 (Figure S4). However, this value (α=1) fails when used to determine the distribution of the samples with a %CV equal to 5 % due to an over-smoothing effect (Figure S4).

**Table 1.** Spearman’s correlation between the estimated and true standard deviation for four groups of samples with a constant mean hydrodynamic radius and different standard deviations (eleven subsamples). Correlation plots are shown in Figure S3.

| $R_h$ | $\alpha: 0.0001$ | $\alpha: 0.001$ | $\alpha: 0.01$ | $\alpha: 0.1$ | $\alpha: 1$ |
| --- | --- | --- | --- | --- | --- |
| 2 nm | 0.6 | 0.6 | 0.7 | 0.6 | 0.5 |
| 6 nm | 0.7 | 0.8 | 0.9 | 0.9 | 0.9 |
| 18 nm | 0.8 | 0.9 | 0.9 | 1 | 1 |
| 54 nm | 0.6 | 0.7 | 0.8 | 0.9 | 0.9 |

To improve the accuracy of the sample distribution estimation, we evaluated the efficacy of the L-curve criteria. It has been previously shown that this method yields correct particle size distributions for DLS data of microgel suspensions (Scotti *et al*., 2015). The corresponding heuristic rule consists of plotting the logarithm of the residuals (fidelity term) against the logarithm of the norm of the regularised solution (penalty term) for different values of α and selecting the value of α corresponding to the corner point of the L-shaped curve (Figure S5). Hereby, we achieve a balance between the size of the regularised solution and the accuracy of the fit. To find the corner of the L-curve we used the triangle method proposed by (Castellanos *et al*., 2002). In our simulations, this approach resulted in finding values of α which also gave a significant correlation between the expected and estimated standard deviations. However, the α values were sometimes suboptimal (Figure S4). Nonetheless, this strategy is better than arbitrarily selecting a fixed α as seen for example by the distance between the estimated and true intensity distributions for the α = 10^−4^. In Raynals, it is also possible to explore different values of regularisation terms and export them together with the penalty (||Lx|| from Equation 5) and fidelity terms (residuals). This last feature allows users to eventually explore different parameter selection rules.

### 2.3 Addressing Raynals performance with experimental DLS data

To assess the performance of Raynals on experimental data, we conducted DLS experiments using a plate reader (wavelength of 817 nm and detection angle of 150°) on two extensively characterised proteins: carbonic anhydrase (CA) and bovine serum albumin (BSA) (Figure 5). For CA and BSA the estimated R_h_ were 2.3 ± 0.06 and 3.9 ± 0.08 nm (mean ± standard deviation of the WHM), in complete agreement with previously reported values (Graewert *et al*., 2020; Brownsey *et al*., 2003; Jachimska *et al*., 2008).

**Figure 5.**
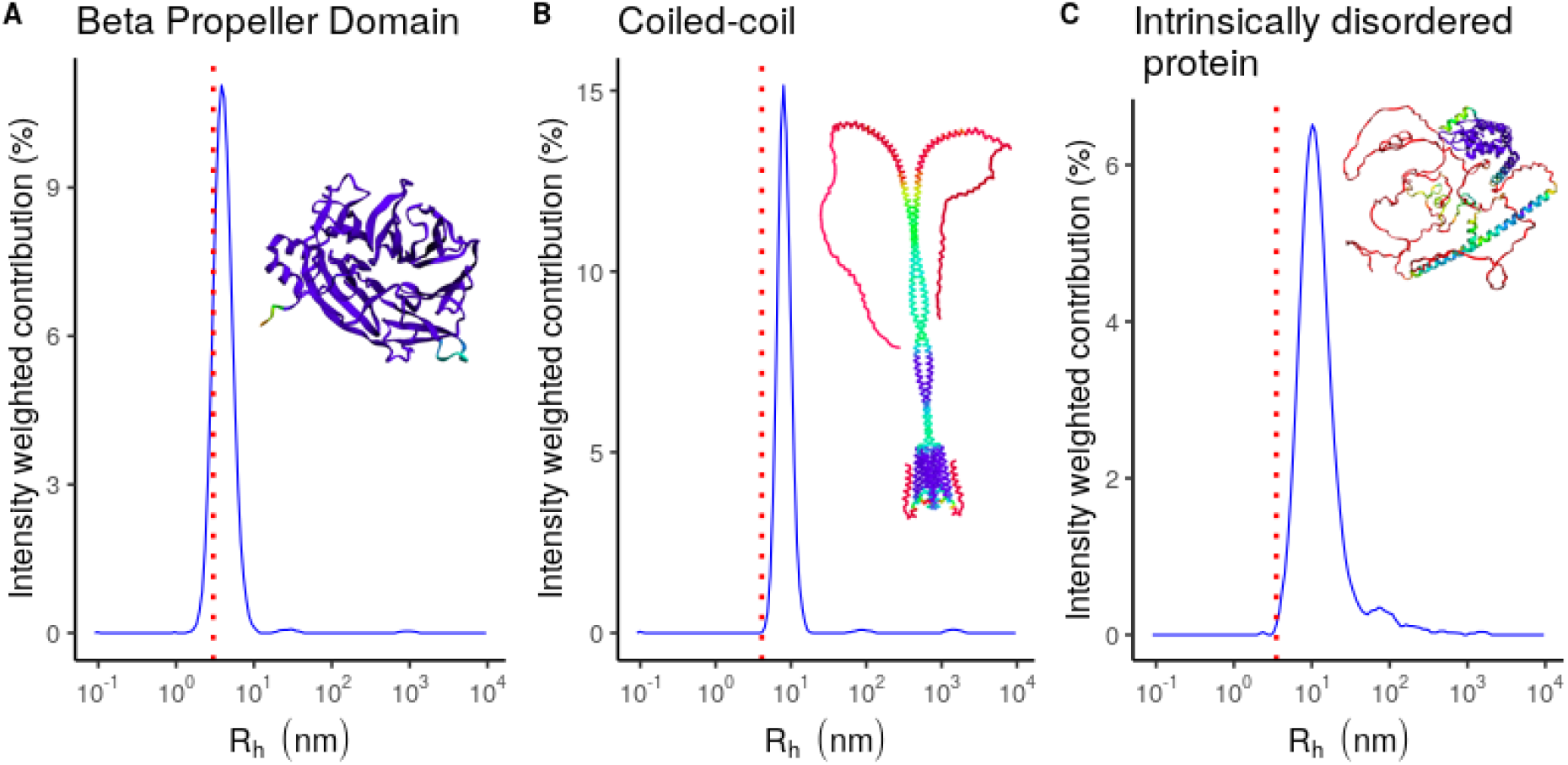
DLS measurements and AlphaFold model predictions of in-house samples. The 30 acquisitions were fitted separately and the estimated intensity values were then averaged. A), B) and C) R_h_ distribution (blue lines) of the Beta-Propeller Domain, Coiled-coil polypeptide and IDP, respectively. Red dashed lines represent the expected R_h_ based on the molecular weight and a globular model. AlphaFold models are coloured by pIDDT values (Very low: red, Low: yellow, OK: green, Confident: Lightblue, Very high: Violet).

In addition, we acquired DLS curves from commercially available gold nanoparticles (GP) with a radius of 2.5, 10 or 20 nm. Since it was not possible to centrifuge the samples due to the sedimentation of the GPs, large dust particles present in the sample heavily contributed to the scattering signal and dominated the measurements for the 2.5 nm GP. For the 10 nm and 20 nm GP sizes, even though the curves were noisy and presented more than one transition, the observed R_h_ values were 11 ± 1.3 nm and 19.5 ± 2.8 nm respectively, in agreement with the expected values (Figure S6).

In addition, we tested the performance of Raynals analysis using a second DLS device with a cuvette holder (wavelength of 658 nm and detection angle of 90°) on in-house produced proteins. We used a beta-propeller domain (BPD), a polypeptide that adopts a coiled-coil structure (CC), and an “intrinsically disordered protein” (IDP). The mean WHM from 30 acquisitions were respectively 3.8 ± 0.2, 7.9 ± 0.4 nm and 10.1 ± 0.6. It’s important to remember that the R_h_ is derived from the diffusion coefficient and requires assuming spherical hard particles (Stokes-Einstein relation, Equation 12).

The BPD is a monomeric globular protein (ter Haar *et al*., 1998) of 40 kDa. The CC has a molecular weight of 53 kDa and is a stable dimer (Yang *et al*., 1999), and the IDP has a molecular weight of 65 kDa. The comparison of the approximated radii for each of these proteins highlights the versatility and limitations of using DLS to assess for example the oligomeric state of the proteins; the globular BPD is not dissimilar in Mr to the IDP although due to the disordered region the calculated R_h_ is almost 2.5 times greater. If the IDP were assessed as a globular protein, the approximate molecular weight would be at least two orders of magnitude different from the correct value. Then, the CC has a radius of two times the value of the BPD although it is only double the weight. For these reasons, absolute R_h_ values should not be used for analysing oligomerization.

### 2.4. Comparison of Raynals to other DLS analysis software

Raynals can be compared with other DLS software such as CONTIN (Provencher, 1982), SEDFIT (Brown *et al*., 2007) and the commercial software DYNAMICS (www.wyatt.com/products/software/dynamics.html) (Table 2). While the last three programs require installation under a certain operating system, Raynals is available online and can be executed from a browser, which facilitates access to potential users independent of the performance of their computer. One of the strong points of CONTIN is the availability of the code as an open source. However, it is written in Fortran, which limits the possibility of users to adapt it for their personal purposes (e.g., changing the regularisation matrix). The code for fitting DLS data from Raynals is available on GitHub (https://github.com/osvalB/dynamicLightScatteringAnalysis) and should be easier to modify, as it was written in Python.

**Table 2.** Comparison of software to analyse DLS data.

| Software | Online | Open source | Open access | Analysis of multiple curves |
| --- | --- | --- | --- | --- |
| Raynals | Yes | Partially* | Yes | Yes |
| CONTIN | No | Yes | Yes | No |
| SEDFIT | No | No | Yes | No |
| DYNAMICS | No | No | No | Yes |

\*We provide the code for the data analysis step (the code for the user interface is not available.).

The four programs use a regularisation approach for data fitting but CONTIN and SEDFIT only allow to fit a single curve at a time. The increase in the availability of plate base DLS, which allows the screening of multiple conditions, creates the need for data comparison tools. Both Raynals and DYNAMICS stand out for their ability to analyse and plot multiple curves at the same time. Raynals is designed with advanced features that allow the comparison of peaks in defined regions of the distribution. Interestingly the analysis of R_h_ is done by automatic peak detection and this is the value that is reported, facilitating the analysis of the region of interest in the distribution. This feature permits easy comparison of R_h_ while the user is experimentally screening for different conditions (eg. different buffers during sample optimisation). Differently, the so far existing software retrieves R_h_ as an average which is more prone to be disturbed by small events or peaks.

Is fair to mention that each software also has unique features. DYNAMICS, for instance, can model concentration-dependent size changes. This is an interesting addition that would allow users to estimate critical micelle concentrations (CMC) of detergents for example. Meanwhile, CONTIN is capable of doing global analysis where an external parameter is varied (e.g. angle of detection) (Provencher & Štêpánek, 1996), SEDFIT allows the use of prior probabilities before fitting the data (e.g., peak location).

Raynals has been designed as a powerful tool to address quality control of biological samples, during sample preparation and optimisation. It possesses useful features for experiment planning and training. Simulation of autocorrelation curves can be done using expected R_h_ from one or many populations of particles. DLS is a simple instrument to use, but data can be easily misinterpreted. The simulation tools available in Raynals allow an understanding of the limitations of the measurement and its resolution, it helps in the detection of aggregates and shows the influence of large particles over the recorded signal. Additionally, Raynals provides flexibility in the way the data is presented: the distribution of decay rates (or diffusion coefficients) can be visualised as histograms, density plots, or grayscale coloured bar plots in publication quality format.

## 3. Conclusions

DLS is a widely used technique to obtain information about the size and dispersity of a macromolecular sample. The fitting of the acquired data entails solving a nonlinear inverse problem, for which the Tikhonov-Phillips regularised inversion has been suggested as a promising approach. Our research, based on both simulations and experimental data, including two DLS instruments, diverse proteins and gold nanoparticles, supports the idea of this method producing reliable results. We look forward to receiving feedback from the scientific community and expanding our software to include more complex analyses such as multi-angle fitting and temperature ramps. Furthermore, Raynals offers a simulation panel that can help users gain a deeper understanding of the challenges of the technique.

## 4. Methods

### 4.1 Input data

The final measurement provided by a DLS experiment is the intensity correlation function (or second-order correlation function). This function, called G_2_(*τ*), can be expressed as an integral over the product of intensities at time *t* and delayed time *t* + *τ*

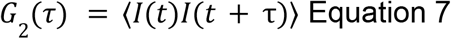

where *τ* is the lag time between two time points. Generally, DLS instruments export the normalised version of G_2_(*τ*):

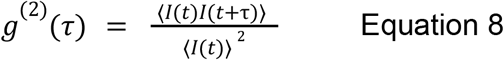

The normalised second-order autocorrelation function g^(2)^(*τ*) is the input data for Raynals.

### 4.2. Fitting DLS data

#### 4.2.1. Theory

The normalised second-order autocorrelation function g^(2)^(*τ*) can be related to the normalised first-order correlation function g_1_(*τ*) through the Siegert Equation (Siegert, 1943):

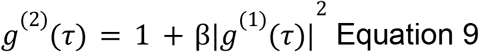

where β is the coherence factor that depends on the instrument and the scattering properties of macromolecules. The function g^(1)^(*τ*) contains information about the motion of the particles and for monodisperse samples, it decays exponentially according to one decay constant. On the other hand, for polydisperse systems, g^(1)^(*τ*) is represented by an intensity-weighted integral over a distribution of decay rates G(┌) (Equation 1). Each decay rate is associated to a certain diffusion coefficient according to the following Equation:

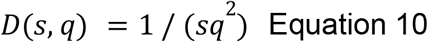

where s is the inverse of the decay rate and q is the Bragg wave vector defined as:

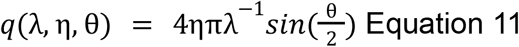

where λ, η, θ are respectively the wavelength of the incident light, the solvent refractive index and the angle of detection. Finally, the diffusion factors (D) can be transformed to hydrodynamic radii (R_h_) with the Stokes-Einstein relation:

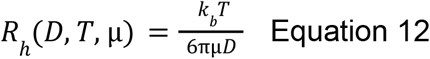

where T and μ are respectively the temperature and viscosity, and *k*_*b*_ is the Boltzmann constant.

##### 4.2.2. Fitting algorithm (Tikhonov-Philips regularised inversion)

Raynals fits the first-order autocorrelation data based on the so-called Tikhonov-Philips regularised inversion. For this purpose, we first obtain β by fitting a second-degree polynomial to the DLS data at times shorter than five μs. Then, we apply Equation 8 to calculate g_1_(*τ*). Due to the square root in this Equation, g_1_(*τ*) can be computed only when g^(2)^(*τ*) ≥ 1. Therefore, we only evaluate the data before the first occurrence of g^(2)^(*τ*) < 1.

After calculating g^(1)^(*τ*), we discretize the decay rate space by using *n* points between 0.1 and 10^6^ nm log spaced in the hydrodynamic radius scale.

The equation we need to fit becomes

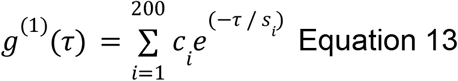

subject to the constraints

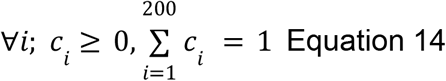

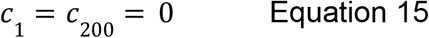

where c_i_ is the i-th contribution of the i-th inverse decay rate (s_i_). Due to the ill-condition nature of the problem (infinite possible solutions), we need to add a regularisation term, so we solve simultaneously the following equations

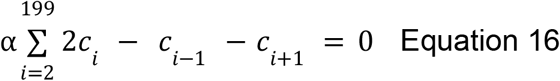

where α is a regularisation parameter controlling how close the relative contribution of each (inverse) decay rate should be to its neighbouring (inverse) decay rates. The whole set of linear equations is solved together using the non-negative least squares solver from the Scipy package (docs.scipy.org/doc/scipy/reference/generated/scipy.optimize.nnls).

#### 4.2.3. Implementation of the L-curve criteria

To build the L-curve, a sequence of regularisation parameters (α) evenly spaced in a log scale are evaluated. This sequence is generated using the following formula:

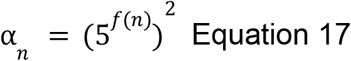

where *f(n)* depends on three parameters called ‘start’, ‘stop’ and ‘step’, and is defined as

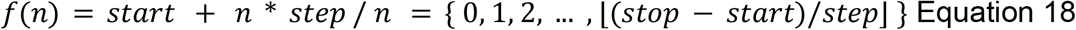

The corner of the curve is then detected by applying the triangle method (Castellanos *et al*., 2002).

### 4.3. Simulated DLS data

#### 4.3.1. Generation protocol

All the artificially generated data is based on the light scattered by populations of particles that have a normal distribution in the hydrodynamic radius space. The necessary steps are:

1. Obtain a sample of particles with a certain normal hydrodynamic radius distribution.
2. Compute the intensity of the scattered light using the Mie Theory (implemented in the Miepython package) by each particle. This step requires selecting an angle of detection, laser wavelength and refractive index.
3. Discretize the hydrodynamic radius space using a log scale from 0.1 to 10^6^ nm and calculate the amount of scattered light in each interval.
4. Divide the scattered light of each interval by the total scattered light to obtain the relative contributions.
5. Convert each hydrodynamic radius into diffusion coefficients at a certain temperature and viscosity.
6. Apply Eq. 12 and then Eq. 8 to calculate the final autocorrelation curve.
7. Add uncorrelated normally distributed error to the autocorrelation curves.

For all the simulations, the wavelength, detection angle, temperature, refractive index and viscosity were set to 817 nm, 150º, 1.33 and 0.00089 Pa s, respectively. The parameter β (from Equation 8) and the standard deviation of the normally distributed error were set to 0.2 and 0.002.

#### 4.3.2 Analysis protocol

For the fitting, the same values of refractive index and viscosity were used. The ‘start’, ‘stop’ and ‘step’ values to build the L curves were respectively -6, 1 and 0.2. The region of interest to estimate the sample R_h_ was 0.1 - 100 nm.

### 4.4. Experimental samples

#### 4.4.1. Commercial samples

Carbonic anhydrase (CA) from bovine erythrocytes, monomer bovine albumin and gold nanoparticles (GNP) were purchased from Sigma Aldrich (CAS 9001-03-0, CAS 9048-46-8, Product number 741949 (radius of 2.5 nm), Product number 741965 (radius of 10 nm) and Product number 741981 (radius of 20 nm). CA and BSA were dissolved in PBS buffer pH 7.4.

#### 4.4.2. In-house produced samples

The recombinant expression of a beta-propeller domain (BPD), intrinsically disordered protein (IDP), and coiled-coil dimeric polypeptide (CC) was done according to the following protocol (protein sequences are provided in the SI). BL21 Gold (DE3) cells containing the pLysS plasmid were transformed with pETM30 plasmids containing the respective cDNA for the globular and coiled-coil polypeptides with an N-terminal fusion of His6 - TEV cleavage site - Glutathione S Transferase (GST) affinity purification tag. Cells were grown in 2xYT media to an OD of 0.6 before induction with 0.2mM IPTG and left shaking at 20 °C overnight. Cultures were harvested at 7,000 rpm x for 20 minutes before resuspension in the lysis buffer. Lysis buffer contains 30mM Tris pH 8, 200mM NaCl, 5% Glycerol. Cells were passed several times through an Emulsiflex and cell debris were sedimented at 38,000 x g for 1 hour at 4 degrees. The proteins were purified using immobilised metal affinity chromatography (IMAC). After the purification tag removal by TEV protease and a second round of IMAC, the target proteins were further purified by gel filtration (Superdex 200 Increase 10/300 GL) in SEC buffer (30 mM Tris pH 8, 150 mM NaCl, 1mM DTT). For the IDP we used the same protocol but with a C terminal His6 tag in a pnEA vector transformed into BL21 DE3 cells without any additional plasmids.

### 4.4. Dynamic light scattering experiments

DLS experiments for CA, BSA & the GP were carried out using a Wyatt DynaPro Plate Reader III instrument (wavelength of 817 nm and angle of detection of 150°). CA and BSA were centrifuged at 17000 rpm for 15’ before measurement (the GP were not centrifuged). DLS experiments for the beta-propeller, IDP and coiled-coil were done using a Wyatt DynaPro NanoStar (cuvette holder, wavelength of 658 nm, and angle of detection of 90°). The globular domain, IDP and coiled-coil were centrifuged at 21000 G for 20’ before measurement. The temperature was set to 20ºC. The running parameters used to obtain the DLS curves are described in Table 3.

**Table 3.** Experimental setup for the DLS experiments.

| Experiment name | Technical replicates | Acquisition time (seconds) | Number of acquisitions |
| --- | --- | --- | --- |
| CA | 5 | 10 | 20* |
| BSA | 5 | 1 | 20* |
| GP | 2 | 1 | 6* |
| BPD | 1 | 5 | 30 |
| IDP | 1 | 5 | 30 |
| CC | 1 | 5 | 30 |

\*Each final DLS curve represents the average of the total number of acquisitions. CA, BSA, beta-propeller, IDP and coiled-coil, were measured respectively at a concentration of 1, 1, 5, 2, 3 (mg/ml). GP (10 nm) and GP (20 nm) were measured using a dilution of 1 to 10 and 1 to 100 from the stock solution.

For the fitting of all samples, the viscosity (0.00089 pascal-second) and refractive index (1.33) of water were used. The ‘start’, ‘stop’ and ‘step’ values to build the L curves were respectively -6, 2 and 0.25.

### 4.5. AlphaFold models

The AlphaFold structure of the BPD, CC and IDP was obtained with the ColabFold v1.5.2: AlphaFold2 using MMseqs2 (Mirdita *et al*., 2022). For the CC, we predicted the homodimer. Default parameters were used for all proteins.

### 4.6. R_h_ prediction

The R_h_ were predicted using the online tool provided by Fluidic Analytics available at https://www.fluidic.com/toolkit/hydrodynamic-radius-converter. The corresponding MW were 40 kDa, 107 kDa, and 65 kDa respectively for the BDP, CC (homodimer), and IPD.

## Figures

Figures 1 and 2 were done with Inkscape (Harrington & Others, 2004). Figure 3 was done with the R package ggplot2 (Wickham, 2016). The plots from Figure 4 were directly exported from Raynals and joined with Inkscape.

## Data and code availability

The experimentally and artificially generated DLS data together with R scripts to produce Figure 3, Figure 5 and Figures S1 to S4 can be downloaded at Zenodo (doi: 10.5281/zenodo.7856850). The Python code to fit DLS data is available at https://github.com/osvalB/dynamicLightScatteringAnalysis.

## Acknowledgments

We acknowledge technical support by the Sample Preparation and Characterisation (SPC) facility at EMBL Hamburg, Germany and the “Molecular Biophysics’’ facility (french acronym: Plate Forme de Biophysique MoleculaIre, PFBMI) at Institut Pasteur, Paris, France. We thank the users of the SPC and PFBMI for constructive feedback. OB is funded by the ARISE fellowship (EMBL and the Marie Skłodowska-Curie Actions). GD-B is funded by the EMBL International Ph.D. programme. OB thanks the EU-funded MOlecular-Scale Biophysics Research Infrastructure (MOSBRI) consortium for his TNA visit (project No MOSBRI-2021-50).

## Author contributions

Project design was done by OB and MGA. Programming of the online tool Raynals was done by OB. DLS experiments of CA, BSA and GP were done by OB and BR. DLS experiments of the globular, intrinsically disordered, and coiled-coil polypeptides were done by GD-B. Resources, funding acquisition, and supervision were carried out by PE and MGA. A first draft was written by OB and MGA and further edited by all authors.

