## Supplementary Information for "Raynals, an online tool for the analysis of dynamic light scattering"

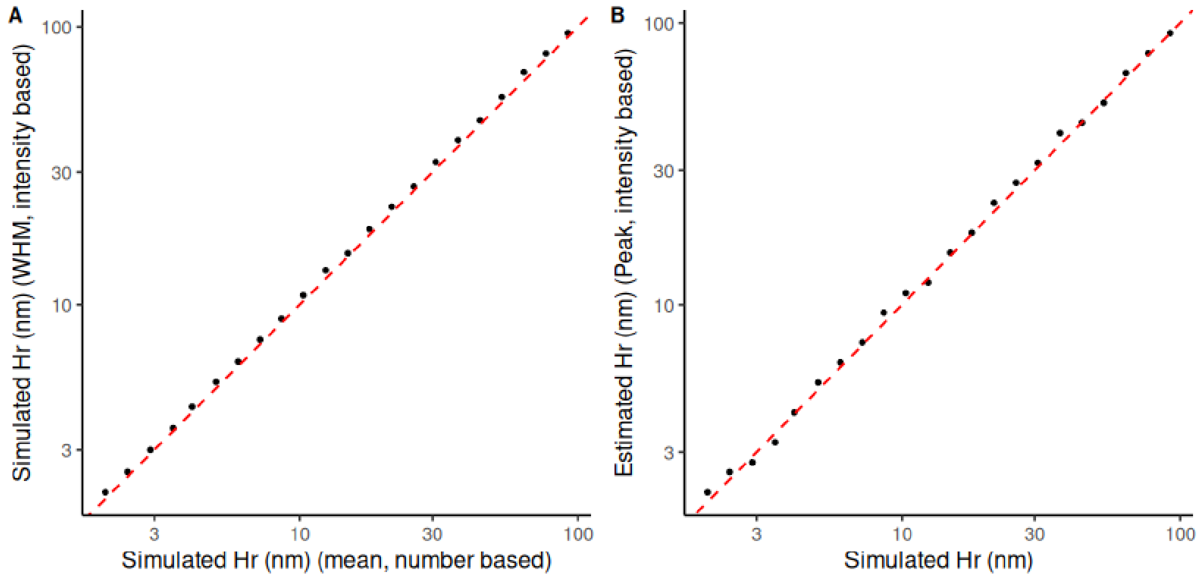


**Figure S1.** A) Mean Hr from the generated number distribution versus WHM from the intensity distribution (transformed from the number distribution based on Mie Theory). B) Estimated Hr based on the mode (peak) versus the Mean Hr from the simulated number distribution. The simulation procedure is described in the Methods Section ‘Case 1 - One population,low CV’.

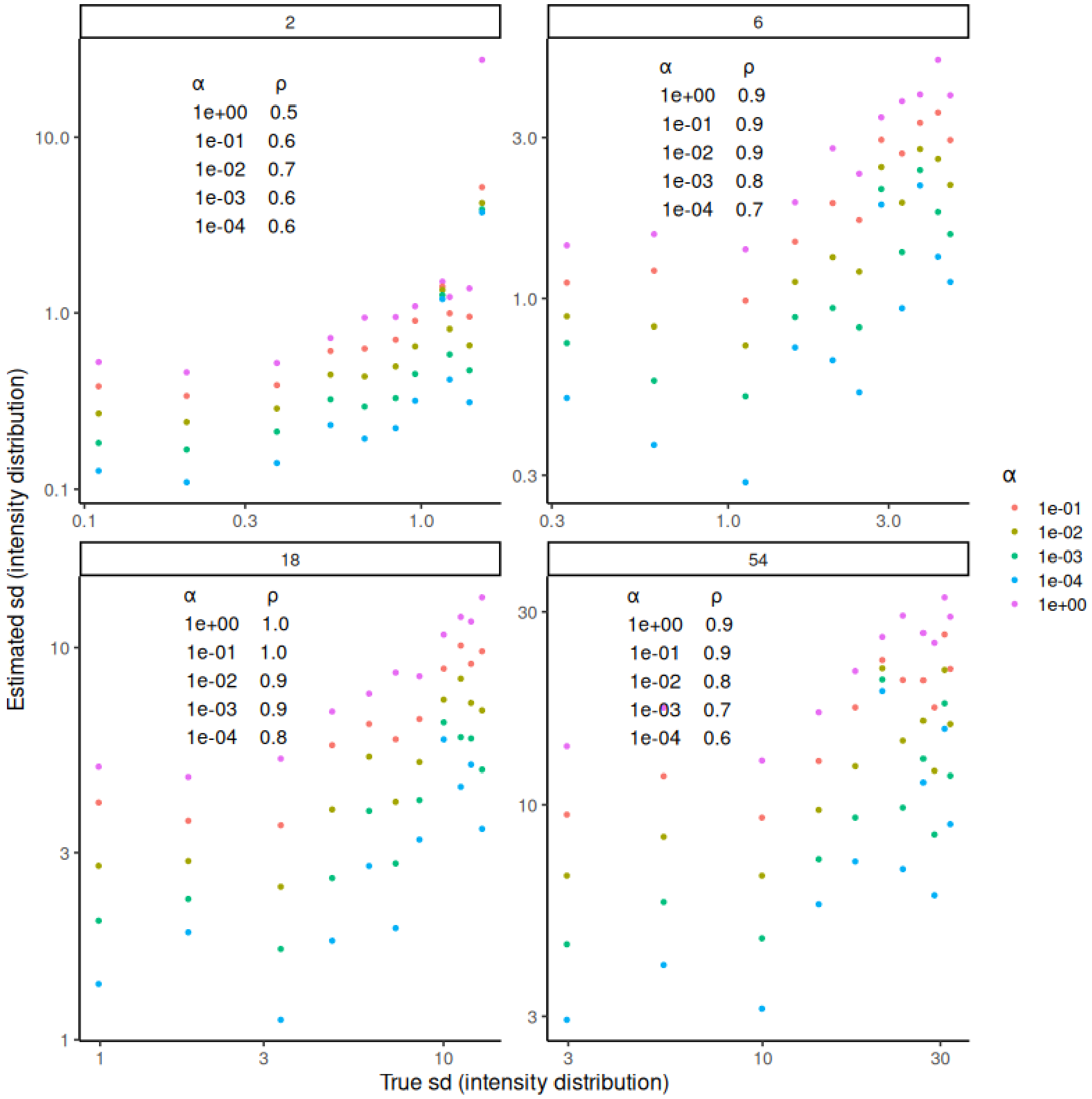


**Figure S2.** Estimated standard deviation (sd) based on the fitted intensity distribution versus standard deviation from the true underlying intensity distribution. Five different values of the regularisation parameter α were tested. Correlation values (ρ) were calculated using Spearman's method. Different panels represent samples with the same mean in the number distribution (2, 6, 18 and 54 nm).

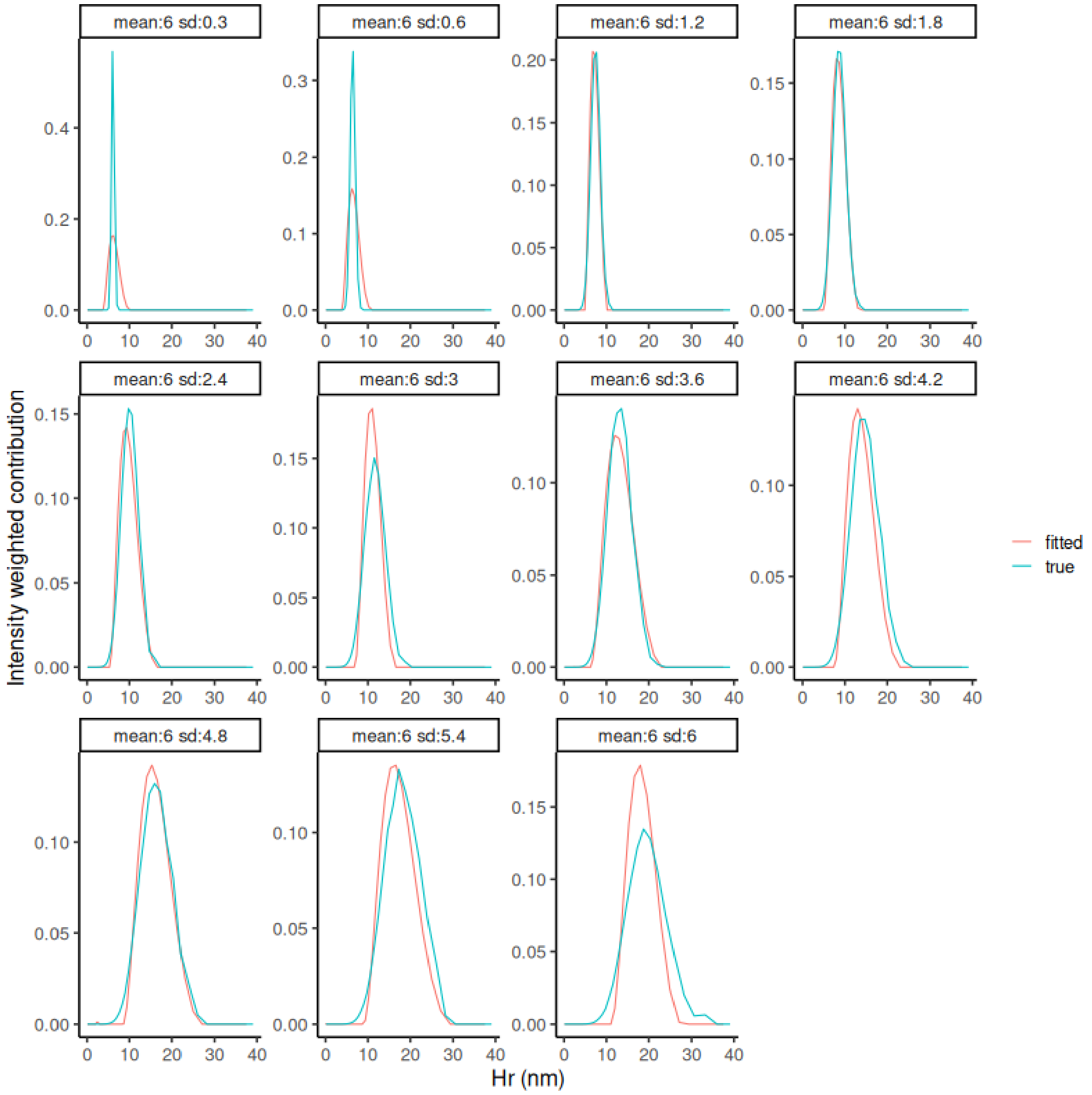


**Figure S3.** Fitted versus true intensity distributions for the simulated samples. Number distributions were first generated with a mean of 6 nm and different standard deviations (0.3, 0.6, 1.2, 1.8, 2.4, 3, 3.6, 4.2, 4.8, 5.4 and 6) and then transformed to intensity distributions. The selected alpha was 0.1.

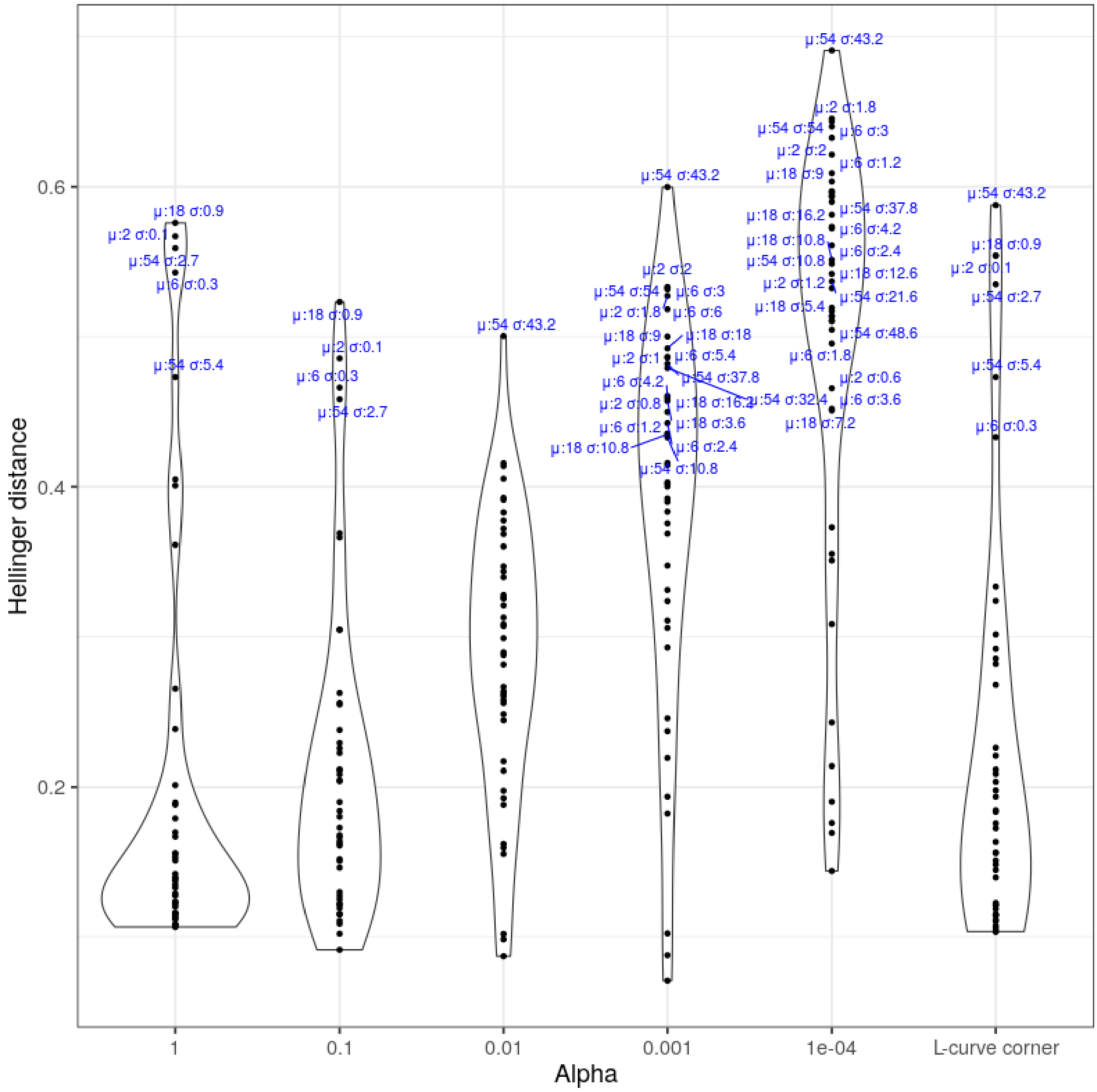


**Figure S4.** Hellinger distance between the estimated and true intensity distributions for the different values of alpha explored. In the case of the L-curve corner, the value of alpha differs for each fitted curve. Labels showing the mean and sd from the original number-weighted distribution were added to all points where the Hellinger distance was larger than 0.42.

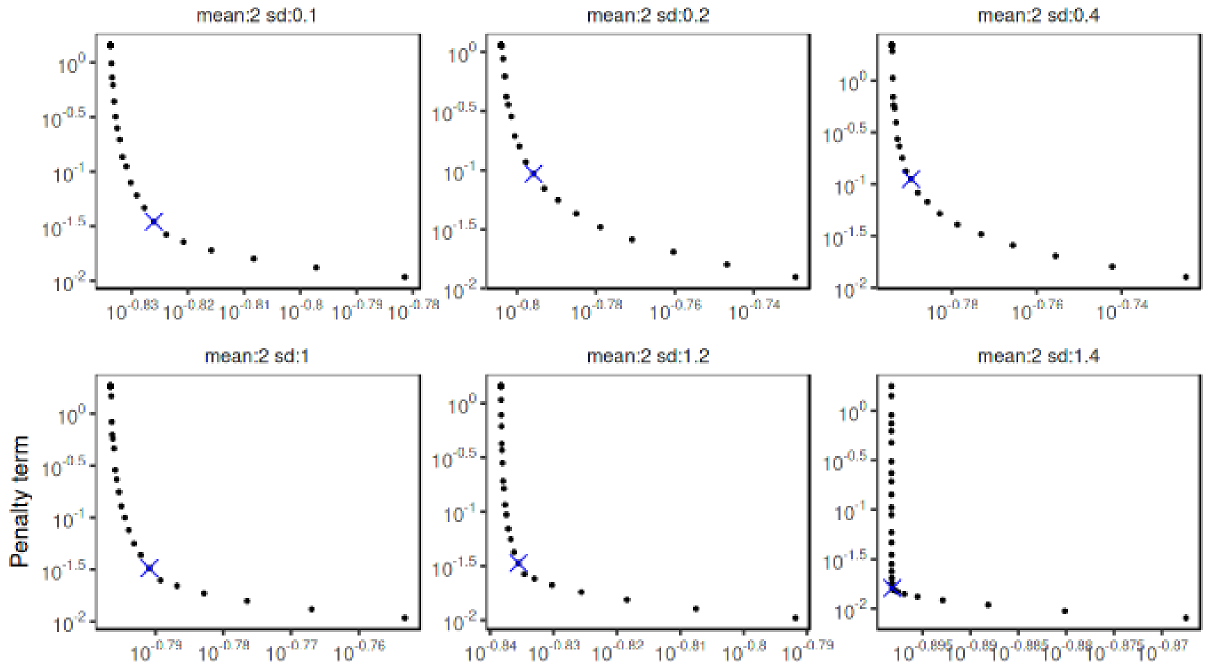


**Figure S5.** Example of log-log scaled curves of the fidelity and penalty term showing an L-shaped form. The blue crosses represent the automatically detected corner. The screenshot was taken from the Raynals app (available at spc.embl-hamburg.de).

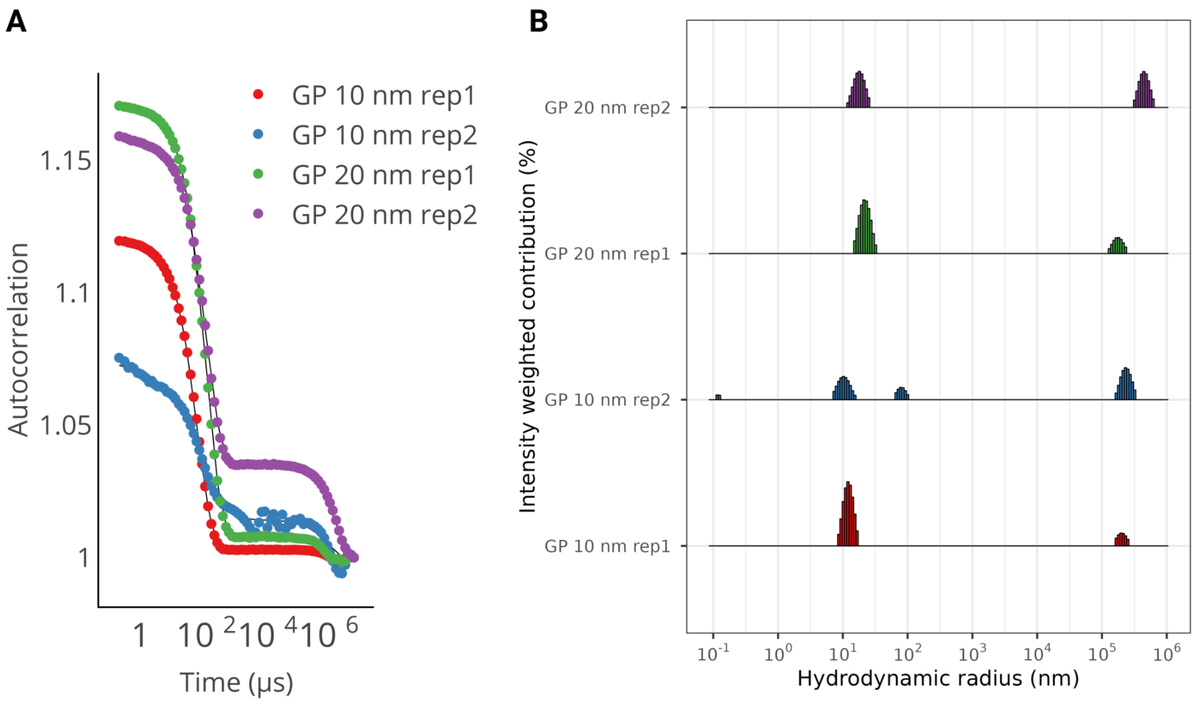


**Figure S6.** A) DLS curves of gold nanoparticles with an hydrodynamic radius of 10 or 20 nm (two technical replicates, where each curve is the average of six acquisitions). B) Estimated hydrodynamic radius distribution.

### Supplementary Methods

#### In-house produced samples - Protein sequences

**Beta-Propeller Domain**
MAPALPIKFQELLQLSSLGVNPSSITFNTCTLESDSFICIRDKKDEVSQPEVLIVDLKNSNNVIRRPIKAD
SAIMHWSRQVIALRAQARTLQIFDLEAKQKLKSTTMSEDVVFWKWVSETTLGLVTEHGIYHWDVFDPT
QAAPVKVFDRNANLQNNQIINYRVSADGKWMVVVGISQQQGRVVGALQLYSKDRGISQAIEGHAAAF
GTIRLDGAPEDTKLFSFAVRTAVGAKLHIVEVDHPETNPVFPKKAVDIFFPPEASNDFPVALQISQKYGII
YLITKYGFIHLYDLETGTCIFMNRISGETIFTACGDKESKGVLGINRKGQVLFVSADENTIVPYVLESHGT
ELALKLASRAGL

**Coiled-coil polypeptide**
APARTPTPTPPVVAEPAISPRPVSQRTTSTPTGYLQTMPTGATTGMMIPTATGAANAIFPQATAQMQP
DFWANQQAQFANEQNRLEQERVQQLQQQQAQQELFQQQLQKAQQDMMNMQLQQQNQHQNDLIA
LTNQYEKDQALLQQYDQRVQQLESEITTMDSTASKQLANKDEQLTALQDQLDVWERKYESLAKLYSQ
LRQEHLNLLPRFKKLQLKVNSAQESIQKKEQLEHKLKQKDLQMAELVKDRDRARLELERSINNAEADS
AAATAAAETMTQDKMNPILDAILESGINTIQESVYNLDSPLSWSGPLTPPTFLLSLLESTSENATEFATSF
NNLIVDGLAHGDQTEVIHCVSDFSTSMATLVTNSKAYAVTTLPQEQSDQILTLVKRCAREAQYFFEDLM
SENLNQVGDEEKTDIVINANVDMQEKLQELSLAIEPLLNIQSVKSNKETNPHSELVATADKIVKSSEHLR
V

**Epsin IDP**
MSKVIRSVKNVTKGYSSVQIKVREATSNDPWGPTGTQMSEIAQLTYGSSTDFYEIMDMLDKRLNDKG
KNWRHVLKALKVMDYCLHEGSELVVTWAKKNIFIIKTLREFQYIDEEGRDVGQNIRVAARELTALIQDE
ERLRAERNDRKMWKNRVNGVEEYAPQHNRDRHDPPRQRHRQYSEEEDLEYRLAIEASKVQEEEDR
KKRESRRQAEEDDDLAKAIKLSKEEEERRRRELEETNAAALFSDTPTQTQQPQFTGFNQGYQQGNA
VDFFANPIDESQLQMQNQMQMQPTGYLNTAYTGFQPQMTGYPNGYVNPFAQQATVFDPYGQQQQV
FQPQATGYNPYFQQQQAQLQVQQQPQLIPDLPEPTLQPGSNNPWANSSSFQSQQALKPTPTGSNN
PFAQRPSTGYKPTTLSTLPEQKTLTSFSSMSNLQASSSSSPFGSNPFSQQSQNQQNQTQSQKPQRQ
MTEHEAKLNALLAQGEGLDTFGNTGQLRIPAQHTAPGVFVNSAGSGLSRLTAETTGNNPFLKPQATG
VPSAPAQLQIPAATGPASMGLNNNPFARAQQQQSPGEAFTQRGNLITF
